# Individual variations in McGurk illusion susceptibility reflect different integration-segregation strategies of audiovisual speech perception

**DOI:** 10.1101/2023.12.15.571270

**Authors:** Chenjie Dong, Zhengye Wang, Ruqin Li, Suiping Wang

## Abstract

The McGurk illusion is a widely used indicator of audiovisual speech integration. In this illusion, incongruent visual articulations biases the auditory speech percepts, resulting in illusory percepts that different from both auditory and visual inputs. Despite its widespread use, its validity for measuring audiovisual integration has been questioned due to substantial individual variations. Classical forced fusion theories propose that variations in McGurk illusion susceptibility reflect differences in multisensory integration ability, whereas Bayesian causal inference (BCI) theory proposes that these variations reflect different audiovisual integration-segregation strategies used according to unisensory accuracy and inferred casual structures of the auditory and visual signals. To test these two proposals, we investigated the relationships between variations in McGurk illusion susceptibility and unisensory accuracy across testing time (Experiment 1, N =161), task type (Experiment 2, N = 88), and stimuli (Experiment 3, N = 37). We found stable negative correlations between McGurk illusion susceptibility and unisensory accuracy both at the group level and single participant level. Participants with weak illusion susceptibility had higher unisensory accuracy than participants with strong illusion susceptibility. Conversely, participants with strong illusion susceptibility were more likely to incorrectly categorize the auditory and visual stimuli as illusory percepts. Moreover, participants with similar unisensory accuracy showed similar McGurk illusion susceptibility. Consistent with the BCI theory, our results indicate that individual variations in McGurk illusion susceptibility reflect different integration-segregation strategies used by participants due to variations in unisensory accuracy. Failing to perceive an illusion does not indicate a failure to integrate audiovisual signals; instead, it indicates a successful segregation of incongruent signals. These findings suggest that caution is necessary when generalizing variations in McGurk illusion susceptibility to differences in audiovisual integration ability.

## 1. Introduction

During face-to-face communication, humans flexibly integrate and segregate acoustic speech signals and visual articulatory signals to improve the detection, categorization, and comprehension of speech information (Holler & Levinson, 2019; Keough et al., 2019; Young et al., 2020). The McGurk illusion is the most compelling example of audiovisual speech integration (Alsius et al., 2018; MacDonald, 2018; McGurk & Macdonald, 1976). In this illusion, physically incongruent auditory and visual signals induce auditory percepts that are different from both the auditory and the visual signals. For instance, when an auditory syllable ba is accompanied by a facial articulation of syllable ga, participants perceive an illusory syllable “da.” However, while it is widely used as an indicator of audiovisual speech integration (Alsius et al., 2018; Tiippana, 2014; Wallace et al., 2020), the McGurk illusion has been questioned as a valid measure of audiovisual integration due to substantial individual variations (Getz & Toscano, 2021; Van Engen et al., 2017, 2022).

The McGurk illusion demonstrates the multisensory characteristics of human speech perception and has been widely used as an indicator of audiovisual speech integration (Alsius et al., 2018; Tiippana, 2014; Wallace et al., 2020). Variations in illusion susceptibility have been used to represent the developmental trajectories of audiovisual integration ability across different age groups both in the healthy population (Burnham & Dodd, 2004; Sekiyama et al., 2014) and in individuals with sensory deficits such as hearing deficits (Schorr et al., 2005; Stropahl et al., 2017) and amblyopia (Narinesingh et al., 2015). Several clinical populations, including individuals with autism (Beker et al., 2018; Feldman et al., 2022; Stevenson et al., 2014; Woynaroski et al., 2013), schizophrenia (De Gelder et al., 2003; Pearl et al., 2009; Roa Romero et al., 2016), and language impairment (Meronen et al., 2013; Norrix et al., 2007), have been shown to exhibit weaker illusion susceptibility compared with control groups, indicating a potential alteration in their audiovisual integration ability (Pulliam et al., 2023; Zhang et al., 2019).

Despite its widespread use, researchers questioned the validity of using the McGurk illusion as a measure of audiovisual integration due to the substantial individual variations in the illusion susceptibility within the healthy population (Getz & Toscano, 2021; Van Engen et al., 2017, 2022). (Getz & Toscano, 2021; Van Engen et al., 2017, 2022). Individual variations in McGurk illusion susceptibility have been documented in many studies of healthy participants (Basu Mallick et al., 2015; Brown et al., 2018; Dong et al., 2023; Magnotti et al., 2015; Nath & Beauchamp, 2012). For some participants, the McGurk illusion susceptibility was close to 100 percent, while for others near zero. Moreover, McGurk illusion susceptibility is not significantly correlated with other audiovisual speech integration measures (Getz & Toscano, 2021; Van Engen et al., 2017, 2022). Therefore, whether McGurk illusion susceptibility truly reflects audiovisual speech integration ability remains a topic of debate (Brancazio & Miller, 2005; Magnotti et al., 2020). This question is particularly relevant given how commonly the McGurk illusion is used by researchers as an indicator of audiovisual integration.

According to the classical forced fusion theories of audiovisual integration, such as maximum likelihood estimation (MLE), variations in McGurk illusion susceptibility could be attributed to differences in the weights assigned to auditory and visual modality during audiovisual integration (Andersen, 2015; Gurler et al., 2015; Strand et al., 2014). During the audiovisual integration, sensory modalities with higher reliability get higher weights than sensory modalities with lower reliability, and the results of the audiovisual integration are shifted to the more reliable modality (Alais & Burr, 2019; Arnold et al., 2010; Bejjanki et al., 2011; Ernst & Banks, 2002). Therefore, in the McGurk illusion (e.g., visual ga dubbed on auditory ba), participants with strong susceptibility to the McGurk illusion (hereinafter referred to as strong illusion participants) give the visual signal higher weights than the auditory signal (Brown et al., 2018), successfully integrate the two signals, and perceive the auditory labial consonant (i.e., ba) as an alveolar consonant “da” due to the bias of the visual velar consonant (i.e., ga). By contrast, participants with weak susceptibility to the McGurk illusion (hereinafter referred to as weak illusion participants) give the visual signal (i.e., ga) lower weight than the auditory signal (i.e., ba), fail to integrate the auditory and visual signals, and consequently fail to perceive the illusion. Accordingly, previous evidence indicated that when presented with the McGurk stimuli, strong illusion participants had higher lip-reading performance (Brown et al., 2018; Strand et al., 2014) and focused more on the mouth area (Gurler et al., 2015) than weak illusion participants. Neuroimaging studies found that strong illusion participants showed higher activation than weak illusion participants at the posterior superior temporal sulcus (pSTS) and classical multisensory areas (Nath & Beauchamp, 2012, 2012; Szycik et al., 2012). In addition, McGurk illusion susceptibility was correlated with activation of pSTS (Benoit et al., 2010; Nath et al., 2011; Nath & Beauchamp, 2012). In brief, the forced fusion theories and corresponding evidence suggest that individual variations in McGurk illusion susceptibility reflect differences in audiovisual integration ability.

Following the forced fusion theories, an intriguing question is why for some people the integration process would give lower weight to the visual modality than the auditory modality, resulting in weak illusion. More to the point, why do some participants barely report the illusion? This observation is inconsistent with the assumptions of forced fusion theories, which maintain that the lack of an illusion is due to low visual reliability or failure to integrate visual signals because these participants were healthy adults with normal hearing and vision, and they showed failure to integrate visual signals on the McGurk stimuli but not on audiovisual congruent signals or other physically incongruent audiovisual signals (e.g., visual ba dubbed on auditory ga). Hence the classical forced fusion theories seem inadequate to address these questions.

In contrast to the forced fusion theories, the Bayesian causal inference (BCI) theory suggests that variations in McGurk illusion susceptibility may not reflect differences in audiovisual integration ability. According to the BCI model, the audiovisual processes involve arbitration between integration and segregation strategies based on unisensory estimates and the inferred causal structure of the auditory and visual signals (Noppeney, 2020, 2021; Shams & Beierholm, 2022). In the McGurk illusion, auditory and visual speech signals from common sources are integrated, and those from separated sources are segregated (Kimmet et al., 2023; Magnotti & Beauchamp, 2017; Meijer & Noppeney, 2023). Therefore, on the one hand, if the participants accurately estimate the auditory signal (i.e., ba) and visual signal (i.e., ga) and infer that those signals belong to separate sources, the optimal operation would be segregating those signals and reporting the estimates of the most dominate modality (i.e., auditory ba) in the speech task. Therefore, on the one hand, if participants accurately estimate the auditory signal (i.e., ba) and visual signal (i.e., ga) and infer that those signals belong to separate sources, the optimal operation would be segregating those signals and reporting the estimates of the dominant modality (i.e., auditory ba) in the speech task. Hence, there is little necessity for integrating two perceptually incongruent signals into an illusory precept, resulting in weak, or even zero, McGurk illusion susceptibility. On the other hand, if participants inaccurately estimate the auditory signal (i.e., ba as “da”) and the visual signal (i.e., ga as “da”) and infer that these signals belong to the common source (Kimmet et al., 2023), the optimal operation would be integrating these signals based on their relative reliabilities, as proposed by the MLE model , and reporting the most possible and reliable percept (i.e., illusion “da”).

In brief, based on the BCI theory, we speculate that due to variations in unisensory accuracy, variations in McGurk illusion susceptibility reflect differences in the inferred causal structures of the auditory and visual signals and in integration-segregation strategies, rather than differences in audiovisual integration ability. Therefore, we propose two hypotheses regarding the mechanisms underlying the individual variations of McGurk illusion susceptibility. First, McGurk illusion susceptibility and unisensory accuracy will be negatively correlated with each other. Accordingly, participants with weak illusion susceptibility will have higher unisensory accuracy than participants with strong illusion susceptibility. Higher unisensory accuracy will enable participants with weak illusion to infer that the auditory and visual signals belong to separate sources and should not be integrated. Conversely, lower unisensory accuracy will enable participants with strong illusion to infer that the auditory and visual signals belong to a common source and should be integrated. Second, the relationship between McGurk illusion susceptibility and unisensory accuracy will be stable across testing time, task type, and speaker due to the stability of suspensory phoneme categorization abilities in healthy adults.

We carried out three experiments to test these hypotheses (Figure 1). Hypothesis 1 was tested in all three experiments. In Experiment 1A (N =161, Figure 1A), first, we validated individual variations in McGurk illusion susceptibility. Second, we examined the correlation between illusion susceptibility and unisensory accuracy. Third, we compared participants with weak vs. strong illusion susceptibility on their responses on unisensory stimuli and McGurk stimuli. In Experiment 2 (N = 88, Figure 1B), we replicated the results of Experiment 1A with different participants and stimuli. In Experiment 3 (N = 37, Figure 1C), we replicated the results of Experiment 1A and Experiment 2 with different participants and stimuli at single participant level. To test the Hypothesis 2, we manipulated the number of testing times (Experiment 1B), task type (Experiment 2), and speaker (Experiment 3) to investigate the stability of relationships between illusion susceptibility and unisensory accuracy. Specifically, in Experiment 1, we examined the stability of relationships between illusion susceptibility and unisensory accuracy across time (Experiment 1B, N = 65) using syllable categorization tasks. In Experiment 2 (N = 88), we assessed the stability of the relationships between illusion susceptibility and unisensory accuracy across the categorization tasks and detection task. In Experiment 3 (N = 37), using 20 McGurk stimuli, we investigated the relationship between illusion susceptibility and unisensory accuracy across speakers at the single participant and single stimulus levels.

**Figure 1.**
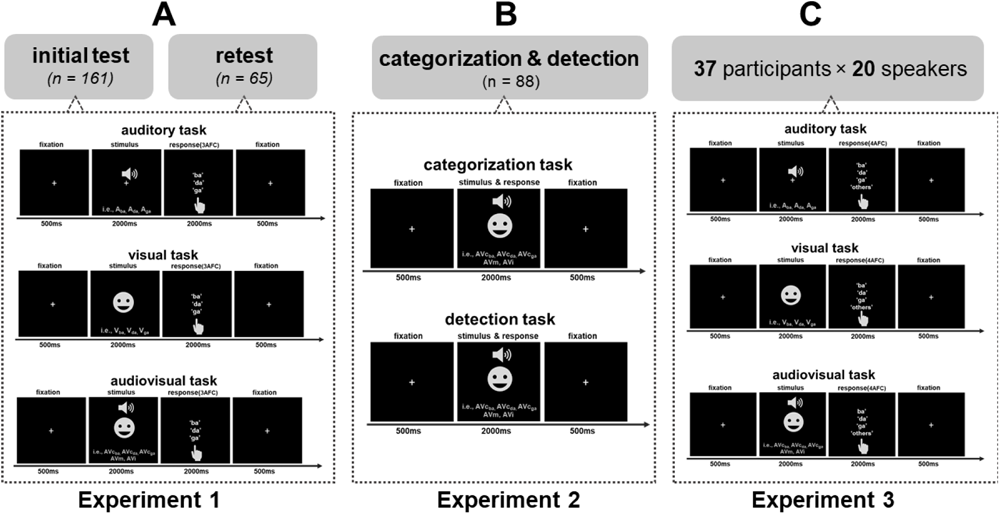
Experimental procedure and stimuli. (A) Auditory, visual, and audiovisual syllable categorization tasks in Experiment 1A and Experiment 1B. (B) Auditory, visual, and audiovisual syllable categorization tasks and the syllable detection task in Experiment 2. (C) Auditory, visual, and audiovisual syllable categorization tasks in Experiment 3. Abbreviations: A_ba_ = auditory ba, A_da_ = auditory da, A_ga_ = auditory ga, V_ba_ = visual ba, V_da_ = visual da, V_ga_ = visual ga, AVc_ba_ = audiovisual congruent ba, AVc_da_ = audiovisual congruent da, AVc_ga_ = audiovisual congruent ga, AVm = McGurk (i.e., auditory ba with visual ga), and AVi = audiovisual incongruent (i.e., auditory ga with visual ba).

## 2. Experiment 1

The purposes of Experiment 1 were to investigate the relationships between McGurk illusion susceptibility and unisensory accuracy and assess the stability of these relationships across testing time. Participants completed the same unisensory and audiovisual syllable categorization tasks in both Experiment 1A (the initial tests) and Experiment 1B (two weeks after the initial tests).

### 2.1 Experiment 1A

#### 2.1.1 Methods

##### Participants

One hundred and sixty-one participants (85 female, age range: 18 - 25) from South China Normal University participated in the Experiment 1A. All participants had normal or corrected-to-normal vision, no history of neurological or psychiatric disorders, and provided informed written consent before the experiment. The protocol for the study was approved by the Human Research Ethics Committee of the School of Psychology at South China Normal University. One participant was excluded because of being unable to follow the instructions; sixteen participants were excluded because their response times were outside the three standard deviations compared with the mean reaction time of all participants.

##### Stimuli

Short clips recorded by two male speakers and two female speakers from previous studies (Basu Mallick et al., 2015) were presented to the participants. For each speaker, there were three auditory stimuli (A: Aba, Ada, Aga), three visual stimuli (V: Vba, Vda, Vga), three audiovisual congruent stimuli (AVc: AVcba, AVcda, AVcga), one McGurk stimulus (AVm = Aba + Vga), and one audiovisual incongruent stimulus (Avi = Aga + Vba). The auditory stimuli were generated by removing the video of the audiovisual congruent stimuli; the visual stimuli were generated by removing the audio of the audiovisual congruent stimuli; the McGurk stimuli were generated by dubbing an auditory ba over the video of facial articulations of ga; the audiovisual incongruent stimuli were generated by dubbing an auditory ga over the video of facial articulations of ba in Adobe Premiere.

The video clips had a duration of around 2 seconds and a resolution of 720×560 pixels. The audio stimuli were recorded at a sampling rate of 48 kHz and were presented to the participants at a sound level of 70 dB using noise-attenuating headphones.

##### Experimental design and procedures

Participants were seated in a sound-attenuating chamber about 60 cm away from a screen (1920 × 1080 resolution). In the syllable categorization task, participants were presented with auditory, visual, audiovisual congruent, McGurk, or incongruent stimuli and indicated the syllable they perceived by pressing one of the 3 buttons (“B”, “D”, and “G” on the keyboard) (figure 1). Each trial started with a fixation cross (0.5 s), followed by the A, V, or AV stimulus (2 s), the three-alternative forced choice (3AFC) screen (2 s). To decrease the attention bias toward any sensory modality and response, we used modality-neutral instruction and required the participants to report the syllables they perceived. Following a strict definition of the McGurk illusion (Alsius et al., 2018; Basu Mallick et al., 2015), we only considered the “da” response, i.e., the fused response, as the illusory percept.

The syllable categorization tasks included three tasks. The unisensory stimuli were presented in a randomized order in an auditory session and visual tasks; the audiovisual stimuli were presented in a randomized order in an audiovisual task. The auditory task included 120 trials (i.e., 10 repetitions per syllable per speaker); the visual task included 120 trials (i.e., 10 repetitions per syllable per speaker); the audiovisual task included 120 congruent trials (i.e., 10 repetitions per syllable per speaker), 80 McGurk trails (i.e., 20 repetitions per speaker), and 80 incongruent trials (i.e., 20 repetitions per speaker). The total duration of the experiment was around 40 minutes.

##### Data analysis

All data were analyzed in the JASP toolbox (https://jasp-stats.org) based on R. As previous studies showed that the distribution of the McGurk illusion susceptibility violated the normal distribution assumption of the parametric statistical tests (Basu Mallick et al., 2015), nonparametric statistical tests were used in the data analysis. For the unisensory and audiovisual congruent stimuli, we calculated the syllable categorization accuracy. For the McGurk stimuli and audiovisual incongruent stimuli, we cannot calculate the categorization accuracy because we used the modality-neutral task. Therefore, we calculated the fractions of each response and compared the fractions of three responses (“ba”, “da”, and “ga”) with the Kruskal-Wallis test to determine which responses were dominant on these stimuli.

##### Responses on the McGurk stimuli

To determine whether the McGurk stimuli induced the illusory percept (“da”), we compared the fractions of illusory response on the McGurk stimuli with the fractions of illusory response on the auditory ba stimuli, the fractions of illusory response on the audiovisual congruent ba stimuli using Kruskal-Wallis test with Conover’s post hoc test.

##### Correlation between the McGurk illusion susceptibility and unisensory accuracy

To assess whether the susceptibility to the McGurk illusion was associated with the unisensory accuracy, we calculated the Spearman correlation coefficients between the fractions of McGurk illusory response and the unisensory categorization accuracy on the auditory counterpart (i.e., auditory ba stimuli) and visual counterpart (i.e., visual ga stimuli) of the McGurk stimuli.

##### Responses on unisensory stimuli in participants with different McGurk illusion susceptibility

To test whether the participants with weak McGurk illusion susceptibility had higher unisensory accuracy than the participants with strong McGurk illusion susceptibility. First, we separated the participants into three groups, which were a group with weak illusion susceptibility (hereinafter referred to as the weak illusion group, McGurk illusion < 0.1), a group with middle illusion susceptibility (hereinafter referred to as the middle illusion group, 0.45 < McGurk illusion < 0.55), and a group with strong illusion susceptibility (hereinafter referred to as the high illusion group, McGurk illusion > 0.9). Then, we compared the unisensory categorization accuracy on the auditory counterpart (auditory ba stimuli) and visual counterpart (visual ga stimuli) of the McGurk stimuli among three groups using the Kruskal-Wallis test. Furthermore, to test whether the strong illusion group more easily miscategorized the unisensory counterparts of the McGurk stimuli as “da” than the weak illusion group, we compared the fractions of “da” response on the auditory ba stimuli and the fractions of ‘da’ response on the visual ga stimuli among three groups using Kruskal-Wallis test.

To examine whether the illusory percept on the McGurk stimuli was fully attributed to the inaccurate “da” percepts on unisensory stimuli, we compared the fractions of “da” responses on McGurk stimuli with the sum of the fractions of “da” responses on unisensory stimuli (auditory ba and visual ga) in the strong illusion group and middle group.

Moreover, to assess whether participants with different illusion susceptibility always assigned different weights to the auditory and visual signals, we compared the fractions of dominant response on the audiovisual incongruent stimuli among three groups using the Kruskal-Wallis test. The assumptions behind these comparisons were that if the individual variations of the McGurk illusion susceptibility are associated with different weights assigned to unisensory signals, these differences should also be shown in the dominant responses to the audiovisual incongruent stimuli.

##### 2.1.2 Results

### Illusory susceptibility and unisensory accuracy

The illusory “da” responses on the McGurk stimuli (mean = 0.54, SD = 0.34, χ^2^ = 189.60, *pbonf* < 0.001, *Kendall’s w = 0.76*) were significantly higher than the auditory ba stimuli (mean = 0.05, SD = 0.09, *pbonf* < 0.001) and the audiovisual congruent ba stimuli (mean = 0.02, SD = 0.05, *pbonf* < 0.001), indicating that the McGurk stimuli induced the illusory “da” percept.

On the McGurk stimuli, the main effect of response was significant (χ^2^ = 85.49, *pbonf* < 0.001, *Kendall’s w = 0.28,* figure 2A). The illusory “da” response (mean = 0.53, SD = 0.34) was significantly higher than the “ba” response (auditory component of the McGurk stimuli, mean = 0.36, SD = 0.35, *T-stat =* 3.65, *pbonf* < 0.001) which was also higher than the “ga” response (visual component of the McGurk stimuli, mean = 0.11, SD = 0.17, *T-stat* = 5.53, *pbonf* < 0.001). Moreover, the susceptibility to the McGurk illusion showed large individual variation (from 0 to 100%, figure 2B) and was negatively correlated with the categorization accuracy of auditory ba (r = -0.36, *p* < 0.001) and visual ga stimuli (r = -0.33, *p* < 0.001, figure 2C).

**Figure 2.**
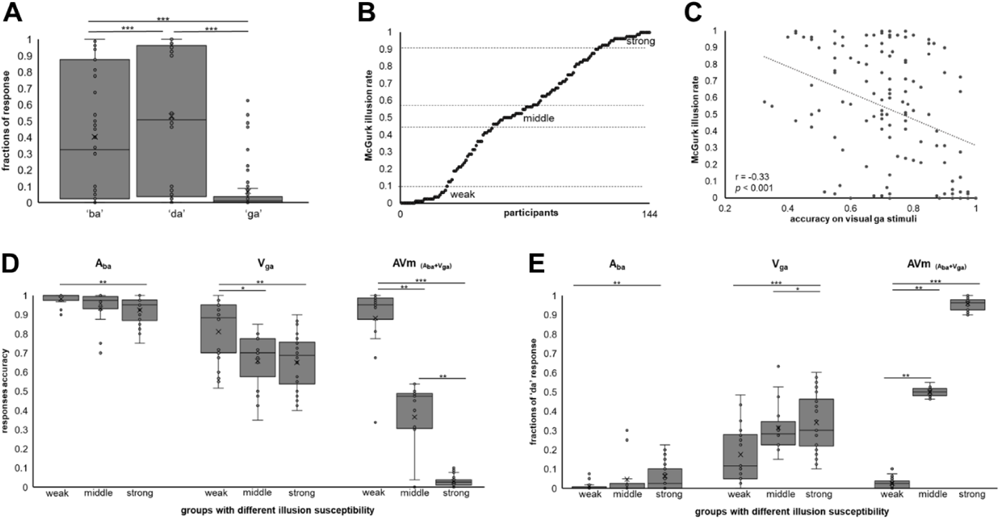
Main results of Experiment 1A. (A) Fractions of responses on the McGurk stimuli. (B) Individual variations of the McGurk illusion susceptibility, each point represents one participant. (C) Correlation between the McGurk illusion susceptibility and visual categorization accuracy. (D) Categorization accuracy on the unisensory counterpart of the McGurk stimuli (auditory ba and visual ga) and the fractions of auditory “ba” response on McGurk stimuli in the weak, middle, and strong illusion groups. (E) Fractions of illusory “da” response on the auditory ba, visual ga, and McGurk stimuli in the weak, middle, and strong illusion groups.

We separated the participants into three groups, which were the weak illusion group (n = 23, mean illusion susceptibility = 0.02, SD = 0.03), middle illusion group (n = 15, mean illusion susceptibility = 0.50, SD = 0.03), and strong illusion group (n = 28, mean illusion susceptibility = 0.96, SD = 0.03) for further analyses.

#### Higher unisensory accuracy in weak illusion group

The unisensory categorization accuracy on the auditory ba and visual ga stimuli was different among the three groups (figure 2D). For the auditory ba stimuli (figure 2D **- Aba)**, the accuracy of the weak illusion group (mean = 0.98, SD = 0.03) was higher than the strong illusion group (mean = 0.92, SD = 0.07, z = 3.26, *pbonf* < 0.01) and marginally higher than the middle group (mean = 0.94, SD = 0.09, z = 1.98, *pbonf* = 0.07).

For the visual ga stimuli (figure 2D **- Vga)**, the accuracy of the weak illusion group (mean = 0.81, SD = 0.16) was higher than the strong illusion group (mean = 0.65, SD = 0.15, z = 3.30, *pbonf* < 0.01) and the middle group (mean = 0.66, SD = 0.15, z = 2.64, *pbonf* < 0.05).

When the visual ga was dubbed on the auditory ba (figure 2D **- AVm**), the weak illusion group reported the auditory component of the McGurk stimuli (mean = 0.88, SD = 0.19) more often than the middle group (mean = 0.36, SD = 0.15, z = 2.77, *pbonf* < 0.01) and strong illusion group (mean = 0.03, SD = 0.03, z = 6.97, *pbonf* < 0.001).

#### Higher illusory “da” response in strong illusion group

The fractions of the “da” response on the auditory ba and visual ga stimuli were different among the three groups (figure 2E). For the auditory ba stimuli (figure 2E **- Aba**), the fraction of “da” response in the strong illusion group (mean = 0.06, SD = 0.07) was higher than in the weak illusion group (mean = 0.009, SD = 0.02, z = 3.24, *pbonf* < 0.01).

For the visual ga stimuli (figure 2E **- Vga**), the fraction of “da” response in the strong illusion group (mean = 0.34, SD = 0.153, z = 3.58, *pbonf* < 0.001) and the middle illusion group (mean = 0.31, SD = 0.13, z = 2.54, *pbonf* < 0.05) were higher than the weak illusion group (mean = 0.17, SD = 0.14). When the visual ga was dubbed on the auditory ba (figure 2E **– AVm)**, the middle group (mean = 0.50, SD = 0.03, z = 2.98, *pbonf* < 0.01) and strong illusion group (mean = 0.96, SD = 0.03, z = 7.51, *pbonf* < 0.001) reported more illusory “da” response than the weak illusion group (mean 0.02, SD = 0.03).

The fraction of “da” responses on the McGurk stimuli was higher than the sum of the fractions of “da” responses on unisensory stimuli (i.e., auditory ba and visual ga) in the strong illusion group (mean = 0.40, SD= 0.17, *p*<0.001, *rbc* = 1) and middle group (mean = 0.36, SD= 0.18, *p*<0.05, *rbc* = 0.72), indicating that the illusion was not fully because of miscategorization on the unisensory stimuli.

#### No group difference on the audiovisual incongruent stimuli

For the audiovisual incongruent stimuli, the main effect of response was significant (χ^2^ = 175.80, *pbonf* < 0.001, *Kendall’s w = 0.61*). Post-hoc analysis showed that the “ga” response (auditory component of the incongruent stimuli, mean = 0.79, SD = 0.34) was significantly higher than the “ba” response (visual component of the incongruent stimuli, mean = 0.20, SD = 0.34, *T-stat* = 7.98, *pbonf* < 0.001) which was higher than the “da” response (mean = 0.01, SD = 0.02, *T-stat =* 5.18, *pbonf* < 0.001).

The fraction of “ga” response on incongruent stimuli was equivalent among the weak illusion group (mean = 0.90, SD = 0.20), middle illusion group (mean = 0.79 SD = 0.35), and strong illusion group (mean = 0.92, SD = 0.14) (**figure S1A)**, indicating that all groups could fully segregate the incongruent auditory and visual stimuli.

### 2.2 Experiment 1B

#### 2.2.1 Methods

##### Participants

Two weeks after Experiment 1A, sixty-five participants from Experiment 1A completed the same tasks (retest) in the same lab.

##### Stimuli

The stimuli were the same as in Experiment 1A.

##### Experimental design and procedures

The design and procedures were the same as in Experiment 1A.

##### Data analysis

The data analysis procedures were the same as in Experiment 1A. To ensure we had enough sample size in the weak illusion and strong illusion groups, we separated the participants into two groups. The weak illusion group included participants whose illusion susceptibility was lower than 0.15. The strong illusion group included participants whose illusion susceptibility was higher than 0.85. Additionally, to test the stability of the McGurk illusion and unisensory accuracy across time, we calculated the Spearman correlation coefficients of unisensory accuracy and McGurk illusion susceptibility between Experiment 1A and Experiment 1B.

##### 2.2.2 Results

The results of Experiment 1B (Figure S2) were the same as Experiment 1A suggesting that the relationship between the McGurk illusion and unisensory accuracy was stable across time. The susceptibility to the McGurk illusion also showed large individual variation (from 0 to 100%, **figure S2B**) and was negatively correlated with the accuracy on visual ga stimuli (r = -0.37, *p* < 0.01, **figure S2C**) and marginally correlated with the accuracy on auditory ba (r = -0.28, *p* = 0.06). The weak illusion group had more accurate unisensory percepts than the strong illusion group (**figure S2D**). The strong illusion group more easily miscategorized the unisensory stimuli as the illusory “da” than did the weak illusion group **(figure S2E**).

In addition, the categorization accuracy on visual ga stimuli (r = 0.69, *p* < 0.001), accuracy on auditory ba stimuli (r = 0.73, *p* < 0.001), and the McGurk illusion susceptibility (r = 0.87, *p* < 0.001) were highly correlated between the Experiment 1A and the Experiment 1B **(figure S1B)**.

## 3. Experiment 2

Experiment 2 was conducted to replicate the results of Experiment 1 and examine the relationships between McGurk illusion susceptibility and unisensory accuracy across tasks. The participants were requested to complete the unisensory and audiovisual syllable categorization tasks and the audiovisual syllable detection task. First, we replicated the results of Experiment 1. Second, we tested the stability of McGurk illusion susceptibility across tasks. Third, we examined the correlation between McGurk illusion susceptibility in the detection task and unisensory accuracy in the categorization tasks.

### 3.1 Methods

#### 3.1.1 Participants

Eighty-eight participants (30 female, age range: 18 - 25) from Guangdong University of Technology participated in this experiment. All participants had normal or corrected-to-normal vision, no history of neurological or psychiatric disorders, and provided informed written consent before the experiment. The protocol for the study was approved by the Human Research Ethics Committee of the School of Psychology at South China Normal University. Eighty-eight participants completed the categorization tasks; seventy-three participants completed the detection task, and sixty-seven participants completed both tasks. Seven participants were excluded because their response times were outside the three standard deviations compared to the mean reaction time of all participants.

#### 3.1.2 Stimuli

Short clips recorded by two Chinese male speakers and one female speaker were presented to the participants. For each speaker, we generated the same auditory, visual, and audiovisual stimuli as Experiment 1in Adobe Premiere toolbox.

#### 3.1.3 Experimental design and procedures

The syllable categorization tasks were the same as Experiment 1. The auditory task included 72 trials (i.e., 8 repetitions per syllable per speaker); the visual task included 72 trials (i.e., 8 repetitions per syllable per speaker); the audiovisual task included 90 congruent trials (i.e., 10 repetitions per syllable per speaker), 60 McGurk trails (i.e., 20 repetitions per speaker), and 30 incongruent trials (i.e., 10 repetitions per speaker).

In the syllable detection task, participants were presented with audiovisual congruent, McGurk, or incongruent stimuli. Each trial started with a fixation cross (0.5 s), followed by an AV stimulus (2 s). Participants were required to press one button (“D” on the keyboard) once they perceived the syllable “da.” This task included 60 target trials (i.e., audiovisual congruent da syllable and McGurk syllable, 10 repetitions per syllable per speaker) and 45 non-targets trials (i.e., audiovisual congruent ba syllable, audiovisual congruent ga syllable, and audiovisual incongruent syllable, 5 repetitions per syllable per speaker). The total duration of the experiment was around 35 minutes.

#### 3.1.4 Data analysis

For the categorization task, the data analysis procedures were the same as in Experiment 1. To ensure the sample size of the groups with strong and weak illusion susceptibilities, we separated participants into two groups.

#### Stability of McGurk illusion susceptibility across tasks

For the detection task, we calculated the detection accuracy on the audiovisual congruent da stimuli and detection rates on the McGurk stimuli. To examine the stability of McGurk illusion susceptibility across tasks, we calculated the Spearman correlation coefficient between the McGurk illusion susceptibility of two tasks.

#### Stability of the relationships between illusion susceptibility and unisensory accuracy across tasks

To test the stability of the relationships between McGurk illusion susceptibility and unisensory accuracy across tasks. First, we separated the participants into two groups, which were the weak illusion group (McGurk illusion rate < 0.3) and the strong illusion group (McGurk illusion rate > 0.8) according to their responses in the detection task. Then, we compared the unisensory categorization accuracy on the auditory ba stimuli and visual ga stimuli in the categorization task between groups using the Mann - Whitney U test. Furthermore, to test whether the strong illusion group more easily miscategorized the unisensory counterparts of the McGurk stimuli as “da” than the weak illusion group, we compared the fractions of “da” response on the auditory ba stimuli and the fractions of “da” response on the visual ga stimuli in the categorization task between groups using Mann -Whitney U test.

### 3.2 Results

#### 3.2.1 McGurk illusion susceptibility in the categorization task

Although we used different stimuli and participants, the results of the categorization task were similar to the results in Experiment 1A. Our stimuli induced the McGurk illusion with large individual variation (**figure S3A**). The illusory susceptibility was correlated with the unisensory accuracy (**figure S3B**). The weak illusion group had more accurate unisensory percepts than the strong illusion group (**figure S3C**). The strong illusion group more easily miscategorized the unisensory stimuli as illusory “da” than the weak illusion group (**figure S3D**).

#### 3.2.2 McGurk illusion susceptibility in the detection task

The McGurk illusion susceptibility in the detection task (mean = 0.51, SD = 0.32) was equivalent to and highly correlated with the illusion susceptibility in the categorization task (mean = 0.52, SD = 0.32, r = 0.82, *p* < 0.001, Figure 3A). The illusion susceptibility in the detection task was marginally correlated with the accuracy of auditory ba stimuli in the categorization task (r = -0.23, *p* = 0.06).

**Figure 3.**
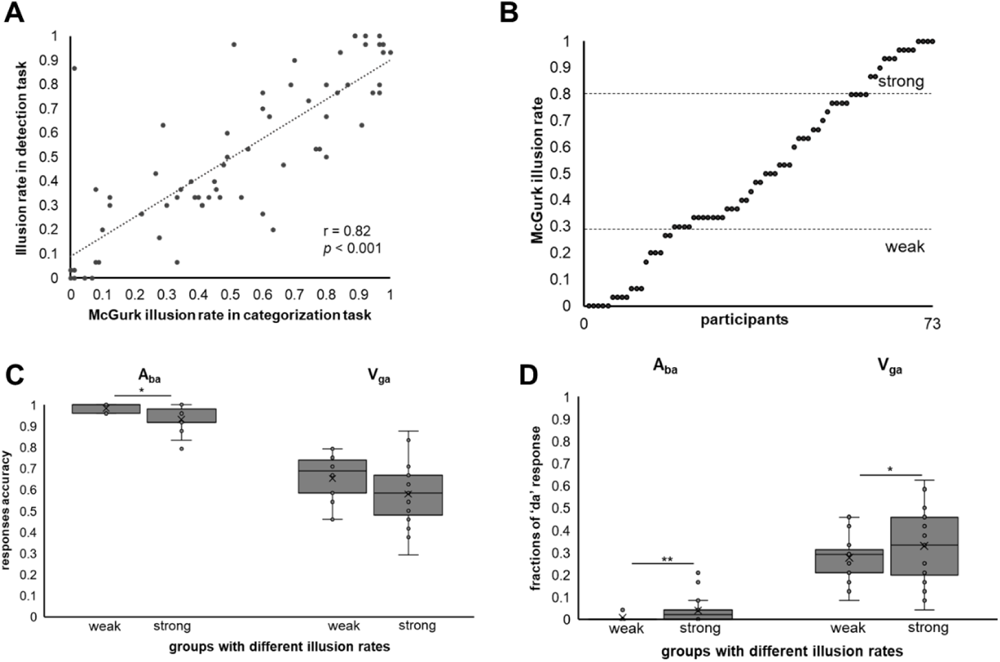
Main results of Experiment 2. (A) correlation between the McGurk illusion susceptibility in the categorization task and detection task. (B) Individual variations of the McGurk illusion susceptibility in the detection task, each point represents one participant. (C) Categorization accuracy on the unisensory counterpart (auditory ba and visual ga) of the McGurk stimuli in the weak illusion group and strong illusion group. (D) Fractions of illusory “da” response on the auditory ba and visual ga stimuli in the weak illusion group and strong illusion group.

We separated the participants into two groups (Figure 3B), according to their illusion susceptibility in the detection task, which were the weak illusion group (n = 14, mean illusion susceptibility = 0.09, SD = 0.10) and the strong illusion group (n = 15, mean illusion susceptibility = 0.94, SD = 0.07) for further analyses.

#### 3.3.3 Higher cross-task unisensory accuracy in weak illusion group

The unisensory categorization accuracy of the auditory ba and visual ga stimuli was different between the two groups (figure 3C). For the auditory ba stimuli (figure 3C **- Aba)**, the accuracy of the weak illusion group (mean = 0.98, SD = 0.02) was higher than the strong illusion group (mean = 0.93, SD = 0.06, *rank-biserial correlation* = -0.53, *p* < 0.05). For the visual ga stimuli (figure 3C **- Vga)**, the accuracy of the weak illusion group (mean = 0.65, SD = 0.12) was equivalate to the strong illusion group (mean = 0.58, SD = 0.16).

#### 3.2.4 Higher cross-task illusory “da” percepts in strong illusion group

The fraction of “da” response on the auditory ba and visual ga stimuli was different between the two groups (figure 3D). For the auditory ba stimuli (figure 3D **- Aba**), the fraction of “da” response in the strong illusion group (mean = 0.06, SD = 0.06) was higher than the weak illusion group (mean = 0.01, SD = 0.02, *rank-biserial correlation* = 0.52, *p* < 0.01). For the visual ga stimuli (figure 3D **- Vga**), the fraction of “da” response in the strong illusion group (mean = 0.35, SD = 0.15) was higher than the weak illusion group (mean = 0.28, SD = 0.08, *rank-biserial correlation* = 0.49, *p* < 0.05).

## 4. Experiment 3

The purposes of Experiment 3 were to investigate the variations of McGurk illusion susceptibility across speakers and test the relationships between McGurk illusion susceptibility and unisensory accuracy at the single participant and single stimulus level. The participants were presented with unisensory and audiovisual stimuli that were recorded from twenty speakers in syllable categorization tasks. Relationships between McGurk illusion susceptibility and unisensory accuracy were tested both on participants with weakest, middle, and strongest illusions susceptibilities and on stimuli that induced weakest, middle, and strongest illusory percepts.

### 4.1 Methods

#### 4.1.1 Participants

Thirty-six participants (17 female, age range: 18 - 25) from South China Normal University participated in the experiment. All participants had normal or corrected-to-normal vision, no history of neurological or psychiatric disorders, and provided informed written consent before the experiment. The protocol for the study was approved by the Human Research Ethics Committee of the School of Psychology at South China Normal University.

#### 4.1.2 Stimuli

Short clips recorded from twenty speakers (10 males and 10 females), seated in front of a dark black background in a sound-attenuating chamber, were presented to the participants. For each speaker, we generated the same auditory, visual, and audiovisual stimuli as Experiment 1 using the Adobe Premiere toolbox. All clips had a duration of around 2 seconds, a resolution of 960×540 pixels, and were presented at an approximate sound level of 70 dB using earphones.

#### 4.1.3 Experimental design and procedures

In the syllable categorization tasks, participants were presented with auditory, visual, audiovisual congruent, McGurk, or audiovisual incongruent stimuli and indicated the syllable they heard or saw (on the visual stimuli) by pressing one of the four buttons (“B”, “D”, “G”, or “O - other “on the keyboard) (figure 1). Compared with Experiment 1 and Experiment 2, we changed the modality that participants need to report on the audiovisual stimuli for two reasons. First, to ensure that the attention of participants was on the same modality, i.e., the dominant modality of speech. Second, fixing the reporting modality allowed us to calculate the response accuracy for both McGurk stimuli and incongruent stimuli. Each trial started with a fixation cross (0.5s), followed by the auditory, visual, or audiovisual stimulus (2s), a four-alternative forced choice (4AFC) screen (2s). The auditory and visual stimuli were presented in a randomized order in the auditory task and visual task; the AV stimuli were presented in a randomized order in the audiovisual task. The auditory session included 600 trials (i.e., 10 repetitions per syllable per speaker); the visual session included 600 trials (i.e., 10 repetitions per syllable per speaker); the audiovisual session included 600 congruent trials (i.e., 10 repetitions per syllable per speaker), 200 McGurk trails (i.e., 10 repetitions per speaker), and 200 incongruent trials (i.e., 10 repetitions per speaker). The total duration of the experiment was around 180 minutes. Participants completed the experiment on two separate days (90 minutes per day).

#### 4.1.4 Data analysis

##### Group level analysis

For each unisensory and audiovisual congruent stimulus of each speaker, we calculated the syllable categorization accuracy. For each McGurk stimulus and audiovisual incongruent stimulus of each speaker, we calculated the fractions of each response. We calculate the illusion susceptibility for each participant by averaging the fractions of illusory response across all speakers and the illusion susceptibility for each speaker by averaging the fractions of illusory response across all participants.

To determine whether the McGurk stimuli induced the illusory percept (“da”), we compared the average fractions of illusory response on all McGurk stimuli, the average fractions of illusory response to all auditory ba stimuli, and the average fractions of illusory response on all audiovisual congruent ba stimuli using Kruskal-Wallis test.

To assess whether the susceptibility to the McGurk illusion was associated with the unisensory accuracy, we calculated the Spearman correlation coefficients between the fractions of the McGurk illusory response and the unisensory categorization accuracy on the auditory ba and visual ga stimuli. Then we used a linear regression model to assess the contribution of the differences in participants, speakers, auditory reliability, and visual reliabilities to the variation of McGurk illusion susceptibility.

##### Single participant-level analyses

To test whether the participants with weak McGurk illusion susceptibility had higher unisensory accuracy than the participants with strong McGurk illusion susceptibility at the single single-participant level. First, we selected three participants with weakest illusion, moderate illusion, and strongest illusion. Second, we compared the unisensory categorization accuracy on the auditory ba and visual ga stimuli averaged across speakers among three participants using the Kruskal-Wallis test. Third, to test whether the strong illusion participant was more easily miscategorized the unisensory counterparts of the McGurk stimuli as “da” than the weak illusion participant, we compared the fractions of “da” response on the auditory ba stimuli and the fractions of “da” response on the visual ga stimuli averaged across speakers among three participants using Kruskal-Wallis test. Lastly, to examine whether the illusory percept on the McGurk stimuli was fully attributed to the inaccurate “da” percepts on unisensory stimuli, we compared the fractions of “da” responses on McGurk stimuli with the sum of the fractions of “da” responses on unisensory stimuli (auditory ba and visual ga) in the strong illusion participant and middle illusion participant.

To further examine the relationship between McGurk illusion susceptibility and unisensory accuracy, we test whether participants with similar unisensory accuracy would have similar McGurk illusion susceptibility based on the logic of representational similarity analysis (RSA). First, we calculated the paired-wise Spearman correlation coefficient of the fraction of illusory response between participants, which resulted in a similarity matrix of the illusion susceptibility of all participants. Then, we calculated the paired-wise Spearman correlation coefficient of the accuracy on auditory ba stimuli between participants, which resulted in a similarity matrix of the auditory accuracy of all participants. Similarly, we calculated the similarity matrix of the accuracy on the visual ga stimuli of all participants. After that, we calculated the Spearman correlation coefficient between the similarity matrix of the illusion susceptibility and the similarity matrix of the auditory accuracy (or similarity matrix of the visual accuracy).

##### Single speaker-level analyses

To test whether the speakers that induced weak McGurk illusion percepts had higher unisensory accuracy than the speakers that induced strong McGurk illusion percepts at the single speaker level. First, we selected three speakers that induced the weakest illusion, moderate illusion, and strongest illusory percepts. Second, we compared the unisensory categorization accuracy on the auditory ba and visual ga stimuli averaged across participants among three speakers using the Kruskal-Wallis test. Third, to test whether the unisensory counterparts of the McGurk stimuli from the strong illusion speaker were more easily miscategorized as “da” than the unisensory counterparts from the weak illusion speaker, we compared the fractions of “da” response on the auditory ba stimuli and the visual ga stimuli averaged across participants among three speakers using Kruskal-Wallis test. To further test the relationship between McGurk illusion susceptibility and unisensory accuracy at the single speaker level, we used the same RSA analysis at the single speaker level analysis.

### 4.2 Results

#### 4.2.1 Varication in McGurk illusion susceptibility

The McGurk stimuli induced the illusory “da” percept (χ^2^ = 68.48, *pbonf* < 0.001, *Kendall’s w = 0.92*). The illusory “da” responses on the McGurk stimuli (mean = 0.54, SD = 0.23) were significantly higher than the auditory ba stimuli (mean = 0.22, SD = 0.15, *pbonf* < 0.01), which was also higher than the audiovisual congruent ba stimuli (mean = 0.06, SD = 0.05, *pbonf* < 0.001).

On the McGurk stimuli, the main effect of response was significant (χ^2^ = 21.46, *pbonf* < 0.001, *Kendall’s w = 0.29*). The fraction of auditory “da” response (mean = 0.54, SD = 0.23) was significantly higher than the “ba” response (mean = 0.23, SD = 0.24, *T-stat =* 3.65, *pbonf* < 0.001) and the “ga” response (mean = 0.21, SD = 0.15, *T-stat* = 5.53, *pbonf* < 0.001).

The McGurk illusion susceptibility showed large individual variation (from 0 to 100%) and was negatively correlated with the categorization accuracy of auditory ba (r = -0.17, *p* < 0.001) and visual ga stimuli (r = -0.23, *p* < 0.001). Moreover, the linear regression model with accuracy on auditory ba stimuli and accuracy on visual ga stimuli as covariates and participants and speakers as factors accounted for 63% variances of the illusion susceptibility (F = 22.68, *p*< 0.001).

#### 4.2.2 Single participant-level analyses

The McGurk illusion susceptibility showed large individual variation across participants (figure 4A). We selected three participants for further analysis: a participant with weakest illusion susceptibility (P18, mean illusion susceptibility = 0.12, SD = 0.19), a participant with moderate illusion susceptibility (P26, mean illusion susceptibility = 0.51, SD = 0.27), and a participant with strongest illusion susceptibility (P30, mean illusion susceptibility = 0.94, SD = 0.07).

**Figure 4.**
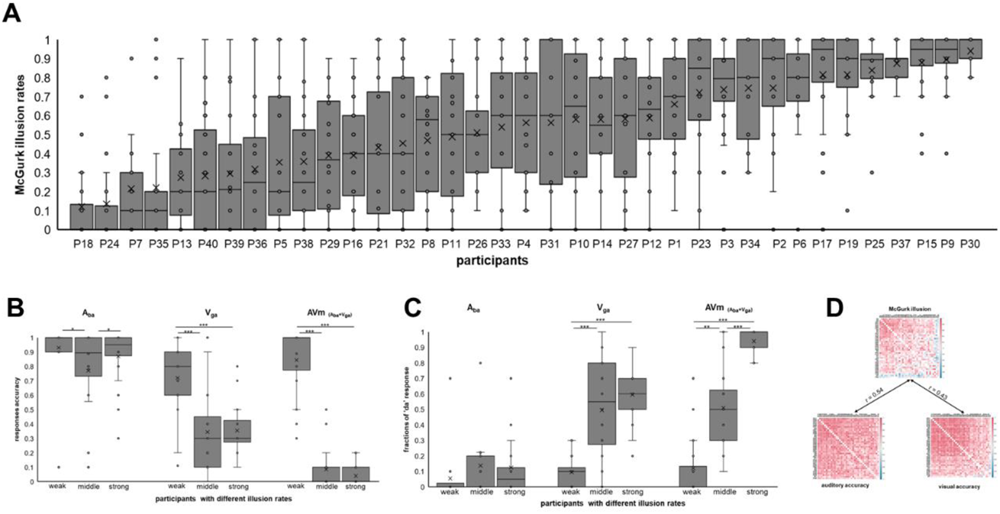
Main results of Experiment 3 – single participant-level analyses. (A) Individual variations of the McGurk illusion susceptibility across participants, each box represents one participant. (B) Categorization accuracy on the unisensory counterpart of McGurk stimuli (auditory ba and visual ga) and the McGurk stimuli in the weakest, moderate, and strongest illusion participants. (C) Fractions of illusory “da” response on the auditory ba, visual ga, and McGurk stimuli in the weakest, moderate, and strongest illusion participants. (D) Similarity matrix of the McGurk illusion susceptibility, auditory accuracy, and visual accuracy across participants.

#### Higher unisensory accuracy in weak illusion participant

The unisensory categorization accuracy on the auditory ba and visual ga stimuli was different among the three participants (figure 4B). For the auditory ba stimuli (figure 4B **- Aba)**, the accuracy of weakest illusion participant (mean = 0.93, SD = 0.2) was equivalent to the strongest illusion participant (mean = 0.87, SD = 0.19) and higher than the moderate illusion participant (mean = 0.77, SD = 0.29, z = 2.66, *pbonf* < 0.05). For the visual ga stimuli (figure 4B **- Vga)**, the accuracy of weakest illusion participant (mean = 0.72, SD = 0.24) was higher than the moderate illusion participant (mean = 0.34, SD = 0.27, z = 3.7, *pbonf* < 0.001) and the strongest illusion participant (mean = 0.35, SD = 0.17, z = 4.05, *pbonf* < 0.001).

When the visual ga was dubbed on the auditory ba (figure 4B **- AVm**), the weakest illusion participant reported the auditory component of the McGurk stimuli (mean = 0.84, SD = 0.23) more often than the moderate illusion participant (mean = 0.08, SD = 0.14, z = 5.28, *pbonf* < 0.001) and the strongest illusion participant (mean = 0.04, SD = 0.07, z = 5.93, *pbonf* < 0.001).

#### Higher illusory “da” response in strong illusion participants

For the auditory ba stimuli (figure 4C **- Aba**), the fraction of “da” response on the strongest illusion participant (mean = 0.12, SD = 0.19) was equivalent to the moderate illusion participant (mean = 0.14, SD = 0.24) and weakest illusion participant (mean = 0.05, SD = 0.16).

For the visual ga stimuli (figure 4C **- Vga**), the fraction of “da” response on the strongest participant (mean = 0.59, SD = 0.16, z = -5.17, *pbonf* < 0.001) and the moderate participant (mean = 0.49, SD = 0.31, z = -4.15, *pbonf* < 0.001) were higher than the weakest participant (mean = 0.09, SD = 0.10).

When the visual ga was dubbed on the auditory ba (figure 4C **– AVm)**, the moderate participant (z = - 3.28, *pbonf* < 0.01) and the strongest participant (z = -6.56, *pbonf* < 0.001) reported more illusory “da” percepts than the weakest illusion participant.

The fraction of “da” responses on McGurk stimuli was significantly higher than the sum of the fraction of “da” responses on unisensory stimuli (mean = 0.72, SD = 0.23, *p*<0.01, *rbc* = 0.82) in the strongest illusion participant.

#### Representational similarity analyses

RSA analyses showed that the similarity matrix of McGurk illusion susceptibility was correlated with the similarity matrix of auditory accuracy (r = 0.54, *p* <0.001) and similarity matrix of visual accuracy (r = 0.42, *p* <0.001) suggesting that participants with similar unisensory accuracy showed similar illusion susceptibility.

#### 4.2.5 Single speaker level analyses

The McGurk illusion susceptibility induced by different speakers showed large individual variation (figure 5A). We selected three speakers for further analysis: a speaker induced weakest illusory percepts (M13, mean illusion susceptibility = 0.22, SD = 0.31), a speaker induced moderate illusory percepts (M1, mean illusion susceptibility = 0.51, SD = 0.33), and a speaker induced strongest illusory percepts (F16, mean illusion susceptibility = 0.90, SD = 0.20).

**Figure 5.**
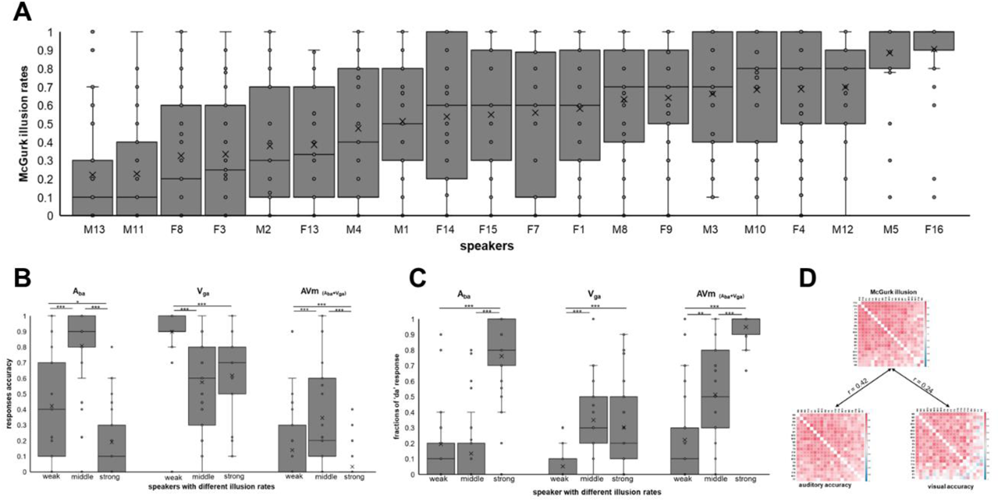
Main results of Experiment 3 - speaker level analyses. (A) Individual variations of the McGurk illusion susceptibility across speakers, each box represents one speaker. (B) Categorization accuracy on the unisensory counterpart of the McGurk stimuli (auditory ba and visual ga) and the McGurk stimuli in the weakest, moderate, and strongest illusion speakers. (C) Fractions of illusory “da” response on the auditory ba, visual ga, and McGurk stimuli in the weakest, moderate, and strongest illusion speakers. (D) Similarity matrix of the McGurk illusion susceptibility, auditory accuracy, and visual accuracy across speakers.

#### Higher unisensory accuracy in weak illusion speaker

The unisensory categorization accuracy of the auditory ba and visual ga stimuli was different among three speakers (figure 5B). For the auditory ba stimuli (figure 5B **- Aba)**, the accuracy on the moderate illusion speaker (mean = 0.81, SD = 0.25) was higher than the weakest illusion speaker (mean = 0.42, SD = 0.35, z = 4.34, *pbonf* < 0.001) which was also higher than the strongest illusion speaker (mean = 0.19, SD = 0.21, z = 2.57, *pbonf* < 0.05). For the visual ga stimuli (figure 5B **- Vga)**, the accuracy on the weakest illusion speaker (mean = 0.89, SD = 0.18) was higher than the strongest illusion speaker (mean = 0.62, SD = 0.29, z = 4.90, *pbonf* < 0.001) and the moderate illusion speaker (mean = 0.57, SD = 0.25, z = 5.83, *pbonf* < 0.001).

When the visual ga was dubbed on the auditory ba (figure 5B **- AVm**), the weakest illusion speaker induced the percept of visual component of the McGurk stimuli (i.e., ga, mean = 0.64, SD = 0.35) more often than the moderate illusion speaker (mean = 0.14, SD = 0.17, z = 5.24, *pbonf* < 0.001) and strongest illusion speaker (mean = 0.06, SD = 0.18, z = 7.62, *pbonf* < 0.001).

#### Higher illusory “da” response in strong illusion speaker

For the auditory ba stimuli (figure 5B **- Aba**), the fraction of “da” response on the strongest illusion speaker (mean = 0.76, SD = 0.24) was higher than the moderate illusion speaker (mean = 0.13, SD = 0.22, z = -7.03, *pbonf* < 0.001) and weakest illusion speaker (mean = 0.19, SD = 0.26, z = -5.91, *pbonf* < 0.001).

For the visual ga stimuli (figure 5B **- Vga**), the fraction of “da” response on the strongest illusion speaker (mean = 0.30, SD = 0.29, z = 4.59, *pbonf* < 0.001) and the moderate illusion speaker (mean = 0.35, SD = 0.22, z = 6.23, *pbonf* < 0.001) were higher than the weakest speaker (mean = 0.05, SD = 0.08).

When the visual ga was dubbed on the auditory ba (figure 5B **– AVm)**, the moderate illusion speaker (z = 2.97, *pbonf* < 0.01) and the strongest illusion speaker (z = 7.60, *pbonf* < 0.001) induced more illusory “da” response than the weakest illusion illusion speaker.

#### Representational similarity analyses

RSA analyses showed that the similarity matrix of McGurk illusion susceptibility was correlated with the similarity matrix of auditory accuracy (r = 0.42, *p* <0.001) and similarity matrix of visual accuracy (r = 0.24, *p* <0.001) suggesting that speakers that induced similar unisensory accuracy showed similar illusion susceptibility.

## 5. Discussion

The cognitive mechanism underlying individual variations in McGurk illusion susceptibility is particularly relevant given that this illusion is widely used as an indicator of audiovisual integration. We examined whether individual variations in McGurk illusion susceptibility reflect differences in audiovisual integration, as suggested by the forced fusion theories, or reflect differences in integration-segregation strategies associated with unisensory accuracy, as proposed by the BCI theory. In three experiments, we found stable correlations between McGurk illusion susceptibility and unisensory accuracy across testing time, tasks, and speakers. Participants with weak illusion susceptibility had higher unisensory accuracy than those with strong illusion susceptibility both at the group level (Experiments 1 and 2) and at the single participant level (Experiment 3). When the weak illusion participants were presented with the McGurk stimuli (i.e., the combination of auditory ba and visual ga stimuli), they could effectively segregate these incongruent stimuli and select what they heard. Conversely, when the strong illusion participants were presented with the McGurk stimuli (i.e., the combination of auditory ba and visual ga), they integrated these noise signals into the illusory “da” percept. Moreover, participants with similar unisensory accuracy showed similar McGurk illusion susceptibility. These results are consistent with the BCI theory (Meijer & Noppeney, 2023; Noppeney, 2021; Shams & Beierholm, 2022). They suggest that individual variations in the McGurk illusion represent different audiovisual integration-segregation strategies that participants use based on unisensory accuracy rather than variations in multisensory integration abilities.

The BCI theory proposed that the ultimate percepts of audiovisual processing are determined by the integration and segregation strategies used by the individuals (Noppeney, 2020, 2021; Shams & Beierholm, 2022). When facing audiovisual speech signals, which are corrupted by internal and external noises, the perceptual system integrates and segregates these signals based on the inferred underlying causal structures (Körding et al., 2007). Audiovisual signals from common sources are integrated by averaging the signals, weighted by relative reliability, whereas those from separate sources are segregated (Magnotti et al., 2020; Magnotti & Beauchamp, 2017). The BCI model can accurately predict human performance in audiovisual spatial localization (Mihalik & Noppeney, 2020; Rohe & Noppeney, 2015), temporal perception (Cao et al., 2019), and speech perception (Magnotti et al., 2020; Magnotti & Beauchamp, 2017; Meijer & Noppeney, 2023).

In accordance with the BCI theory, we found that the participants with different McGurk levels of illusion susceptibility had different unisensory accuracy and used different audiovisual processing strategies. The participants with weak illusion susceptibility had higher unisensory accuracy and estimated the auditory ba stimuli as “ba” percepts and visual ga stimuli as “ga” percepts. Therefore, on the McGurk stimuli, they inferred that the auditory estimation “ba” and the visual estimation “ga” originated from separate sources, correctly segregated them, and reported what they heard (“ba”), resulting in low illusion susceptibility. The participants with strong illusion susceptibility had lower unisensory accuracy and estimated auditory ba signals as “ba” or “da” percepts and visual ga signals as “ga” or “da” percepts (Iqbal et al., 2023; Tiippana et al., 2023). Therefore, on the McGurk stimuli, they inferred that these signals came from common sources and integrated them into the most possible percept (i.e., “da”), resulting in high illusion susceptibility. Recent research also showed that participants inferred a congruent relationship (or common sources) between the auditory and visual signals when perceiving the illusory “da” percept on the McGurk stimuli (Kimmet et al., 2023; Meijer & Noppeney, 2023).

Our findings are also in line with a recent neuroimaging study (Dong et al., 2023) showing that brain activation differences between the McGurk stimuli and audiovisual congruent stimuli could be interpreted by the participants’ perceptual uncertainty, which was related with the sensory accuracy. When participants faced psychically incongruent audiovisual stimuli, those with higher perceptual uncertainty (i.e., lower unisensory accuracy) integrated the noisy auditory and visual signals and reported more illusory percepts than the participants with lower perceptual uncertainty. By contrast, when the perceptual uncertainty on the psychically incongruent audiovisual stimuli was equivalently high, both the low illusion participants and high illusion participants successfully segregated the auditory and visual signals and reported what they heard (**Figure S1A**). Similarly, previous research demonstrated that increased unisensory noise led to higher illusion susceptibility (Stacey et al., 2020), eliminated differences in illusion susceptibility between groups (Ujiie & Wakabayashi, 2022), and enhanced functional connectivity between sensory-specific and unisensory brain areas (Nath & Beauchamp, 2011).

Accordingly, we found a negative correlation between unisensory accuracy and McGurk illusion susceptibility: with an increase of unisensory accuracy, illusion susceptibility decreased (Dong et al., 2023). These results are inconsistent with studies that reported a positive correlation or no correlation between lip-reading abilities and illusion susceptibility (Brown et al., 2018; Strand et al., 2014; Van Engen et al., 2017). The inconsistency may be due to the contents of stimuli used in earlier research and the current research. Earlier research used syllables (Brown et al., 2018; Strand et al., 2014) and sentences (Van Engen et al., 2017) that were different with the counterparts of McGurk stimuli, whereas this study only included the stimuli (auditory ba and visual ga stimuli) that were relevant to the McGurk stimuli. On the other hand, our findings showed that the lower the auditory accuracy, the stronger the McGurk effect, which is consistent with recent studies (Iqbal et al., 2023; Tiippana et al., 2023).

Furthermore, our findings are in line with the observations that individuals with high auditory accuracy, such as musicians, rarely report the McGurk illusion (Proverbio et al., 2016), and individuals with low auditory reliability, such as older adults (Sekiyama et al., 2014) and cochlear implant users (Desai et al., 2008; Stropahl et al., 2017), frequently report the McGurk illusion.

## 6. Conclusion

Taken together, our results suggest that individual variations in McGurk illusion susceptibility are not solely attributable to individual differences in audiovisual integration ability. Instead, these variations appear to be due to different integration-segregation strategies used by participants based on variations in unisensory accuracy. Failure to perceive an illusion does not indicate failure to integrate audiovisual signals; instead, it reflects a successful segregation of incongruent audiovisual signals. Our findings suggest that high caution is required when generalizing the variations in McGurk illusion susceptibility to differences in audiovisual integration ability.

## Data availability statement

Data are publicly available on the Open Science Framework at: https://osf.io/yn9eq/

## Authorship contribution statement

Chenjie Dong, conceptualization, data collection, data analysis, writing manuscript; Zhengye Wang, generating stimuli, data analysis; Ruqin Li, generating stimuli, data analysis; Suiping Wang, conceptualization, resources, writing manuscript, supervision, funding acquisition, project administration. We thank Caizhe Liu and Qiu Yao for helping with data collection.

## Conflict of interest statement

All authors declare that they have no conflicts of interest.

## Acknowledgments

This research was funded by the National Natural Science Foundation of China (No. 32171051).

## Notes

### Competing Interest Statement

The authors have declared no competing interest.

